# Culture and genomic analysis of *Arsenophonus apicola* sp. nov. isolated from the honeybee *Apis mellifera*

**DOI:** 10.1101/2022.01.26.477889

**Authors:** Pol Nadal-Jimenez, Stefanos Siozios, Crystal L. Frost, Rebecca Court, Ewa Chrostek, Georgia C. Drew, Jay D. Evans, David J. Hawthorne, James B. Burritt, Gregory D.D. Hurst

## Abstract

The genus *Arsenophonus* has been traditionally considered to be comprised of heritable bacterial symbionts of arthropods. Recent work has reported a microbe related to the type species *A. nasoniae* as infecting the honey bee, *Apis mellifera*. The association was unusual for members of the genus in that the microbe-host interaction arose through environmental and social exposure rather than vertical transmission. In this study, we describe the *in vitro* culture of ArsBeeUS, a strain of this microbe isolated from *A. mellifera* in the USA. The 16S rRNA sequence of the isolated strain clearly indicates it falls within the genus *Arsenophonus*. Biolog analysis indicates the bacterium has a restricted range of nutrients that support growth. *In vivo* experiments demonstrate the strain proliferates rapidly on injection into *A. mellifera* hosts. We further report the closed genome sequence for the strain. The genome is 3.3 MB and 37.6% GC, which is smaller than *A. nasoniae* but larger than the genomes reported for non-culturable *Arsenophonus* symbionts. The genome is complex, with 6 plasmids and 11 predicted intact phage elements, but notably less complex than *A. nasoniae*. With 92% average nucleotide identity (ANI) to the type strain *A. nasoniae*, the ArsBeeUS strain isolated is clearly distinct from the type species, and we propose the name *Arsenophonus apicola* sp. nov. (CECT 30499; LMG 32504).

## INTRODUCTION

It is now understood that microbes influence multiple aspects of animal biology (1). Symbioses may be formed through the host acquiring a symbiont from the environment, a free-living symbiont infecting a host, or an interplay of host and microbe processes (2). In other cases, symbiosis is a result of vertical transmission, with either the microbe establishing routes of infection to the host germline, the host ‘organising’ symbiont transfer to the germline or transmission to the germ line occurring through the interplay of host and symbiont processes (2, 3). The mode of symbiont acquisition is an important factor shaping the symbiosis, with vertical transmission creating a population bottleneck that both reduces conflict between variants within a host and also aligns transmission success of the microbe to host fecundity (4). Understanding how evolution favours change in transmission mode, and the consequences of it, requires a detailed understanding of clades that retain diversity in this aspect of symbiosis.

The genus *Arsenophonus* is unusual in the diversity of interactions with hosts that are found within the clade. The type species, *Arsenophonus nasoniae*, was identified as the causative agent of the son-killer trait in the parasitic wasp *Nasonia vitripennis* (5, 6). This microbe shows mixed modes of inheritance; vertical transmission is enabled by a pronounced tropism to the host ovipositor (7) while horizontal transmission occurs when two female wasps co-parasitize a fly host (8, 9). The microbe is unusual in its genome complexity, with the main chromosome complemented by 17 plasmids, and these elements combined harbour 55 intact or relic prophage elements (10). Other members of the clade are either facultative heritable symbionts (not-required by the host), such as *Cand.* Arsenophonus triatominarum (11), or symbionts obligately required by the host such as *Cand.* Arsenophonus arthropodicus (12) and *Cand.* Riesia (13). The clade also contains insect-vectored plant pathogenic strains, including *Ca.* Phlomobacter fragariae, the cause of marginal chlorosis of strawberries (14) (15).

Recently, *Arsenophonus* infection has been reported in the honey bee, *Apis mellifera* (16–20). In contrast to the other members of this clade, there was no evidence *Arsenophonus* in *A. mellifera* underwent vertical transmission. Environmental acquisition was the dominant form by which *Arsenophonus* infection occurred, and bees were shown to transmit the microbe horizontally via social interactions (20). There were large seasonal changes in the frequency with which colonies carried *Arsenophonus* infection. Colonies tended to lose infection overwinter, producing a very low frequency of infection in bees on spring emergence, *Arsenophonus* then appeared to be reacquired throughout the summer with the frequency of infected colonies peaking in August/September. Infection with *Arsenophonus* has also been linked to poor bee health (21, 22), however further work is required to corroborate this.

In this paper, we report the *in vitro* culture of this microbe derived from *Apis mellifera* originating in the USA. This data is accompanied by tests of the growth requirement of the bacteria *in vitro* and its capacity to grow *in vivo* in honey bees following injection. We also report completion of the genome sequence for the strain, which we name *Arsenophonus apicola* sp. nov.

## MATERIALS AND METHODS

### Symbiont isolation, morphology *in vitro* and identification through 16S rRNA sequence

Worker *Apis mellifera* from Wisconsin were isolated from a managed colony and hemolymph removed and spread on DNase agar with methyl green (Difco #222020, Voigt Global Distributing, Lawrence KS). Small bacterial colonies appeared after 10 days of growth. 16S rRNA sequence was obtained from PCR product amplified using primers 27F/1492R (23, 24), the product was cleaned and Sanger sequenced using the original primers.

An initial phylogenetic analysis based on the 16S rRNA gene was performed including representative sequences from the database. To this end, a structural alignment of 16S rRNA sequences was generated using the SSU-ALIGN software (25) and purely aligned columns were removed using the ssu-mask command. Phylogenetic relationships were inferred using maximum likelihood (ML) as implemented in IQ-TREE 2.0.3 (26) under the TIM3e+I+G4 model of substitution selected using ModelFinder (27). The robustness of the tree was assessed based on 1000 ultrafast bootstrap replicates (28). The phylogenetic tree was visualized and annotated using the Interactive Tree Of Life (iTOL) server (29).

The strain was preserved at −80°C in 20% glycerol stocks, and then grown routinely on BHI medium. Grams stain status was determined and appearance/size ascertained using scanning electron microscopy (SEM).

### In vitro Growth requirements

Carbon utilization and inhibition by stressful environments/xenobiotics was assessed using BIOLOG GEN III plates (Cat. No. 1030), with the isolated strain compared to the type species of the genus, *A. nasoniae*. 5 ml of both bacterial strains were grown for 48h at 30°C with 250 rpm shaking till their maximum OD600 (0.4-0.6) was reached, the cultures were then spun down, and each pellet was resuspended in a tube of IF-A inoculating fluid (Biolog, Cat. No. 72401) and 100 μl of this suspension was added to each of the 96 wells of the BIOLOG GEN III plate. The plate was subsequently placed to incubate at 30°C without shaking for up to 1 week, to allow *Arsenophonus* spp. growth and full development of the chromogenic medium.

### Capacity to grow in *Apis mellifera*

*Arsenophonus apicola* was transformed with the pOM1-*gfp* plasmid (30) following the method in (7). BHI containing pOM1*-gfp A. apicola* was grown. Concurrently, 690 nl of media were injected, using a Nanoject II (Drummond Scientific), under the 3^rd^ abdominal tergite of each of 32 CO_2_ aneasthetized newly emerged worker *A. mellifera* collected one day previously. This represented an inoculum of 8.2 x 10^5^ cfu, as determined by serial dilution. Control injections were of medium only (22 bees). Bees were removed, maintained in groups of 5-6 at 25 °C in the dark, and then imaged through epifluorescence 1 and 3 days post injection. Bees that died during the experiment were treated likewise but included as a separate category.

### Symbiont Genome sequencing and assembly

The genome of the symbiont was completed using a combination of Illumina and Nanopore reads. High molecular weight DNA was prepared using a phenol-chloroform extraction method (31). Long read Nanopore sequencing was performed using the Rapid Sequencing (SQK-RAD004) protocol (Oxford Nanopore) on a FLO-MIN106 (R.9.4.1) MinION flow cell. Raw Nanopore signals were live basecalled using the processing pipeline implemented in MinKNOW software v18.01.6 (Oxford Nanopore). Illumina sequencing was performed by MicrobesNG (Birmingham) using the Nextera XT library prep protocol on a MiSeq platform (Illumina, San Diego, USA) and reads were adapter trimmed using Trimmomatic 0.30 with a sliding window quality cutoff of Q15 (32). Long read *de novo* assembly was performed with Flye v2.8 using the uneven coverage mode (--meta) and the option --plasmids for recovering short sequences like plasmids. The assembly was further error corrected by mapping the Illumina reads back to it and using six polishing rounds with Pilon v1.23 (33). The raw long reads were mapped back to the assembly and the assembly was visually inspected for mis-assemblies. Genome annotation was performed with the NCBI Prokaryotic Genome Annotation Pipeline (34). Metabolic and functional assessment of the genome was conducted using the Kyoto Encyclopaedia of Genes and Genomes (KEGG) database (35). Prophage regions were predicted using the PHAge Search Tool Enhanced Release (PHASTER) web server (36).

### Phylogenomic and taxonomic classification

The phylogenetic position of the *Arsenophonus apicola* was estimated in relation to 13 publicly available *Arsenophonus* genomes based on the concatenated analysis of 148 single copy core proteins determined using OrthoFinder v2.3.11 (37). Reconstruction methods follow that for the 16S rRNA phylogeny, with the JTT+F+R3 mode selected using ModelFinder (27). The average nucleotide identity (ANI%) values between the *Arsenophonus* genomes were computed using the ANI/AAI-Matrix calculator from the enveomics toolbox collection (38). An additional taxonomic classification of *A. apicola* was performed based on the Genome Taxonomy Database (GTDB) using the GTDB-Tk v1.6.0 software (39, 40) as implemented in the KBase server (41).

## RESULTS

### Symbiont isolation and identification

The bacterium grew optimally at 30 °C in an aerobic environment. Growth was slow, forming colonies after 6 days. Colonies were small (1-2mm), translucent and of a wrinkled appearance on BHI agar. Liquid culture in BHI is similarly slow, with cultures reaching a maximum OD_600_ =0.4-0.6, similar to *A. nasoniae*. The bacterium is gram negative and SEM image reveals a non-flagellated rod-shaped bacterium approx. 2 μm long and 0.5 μm across (Figure 1).

**Figure 1.**
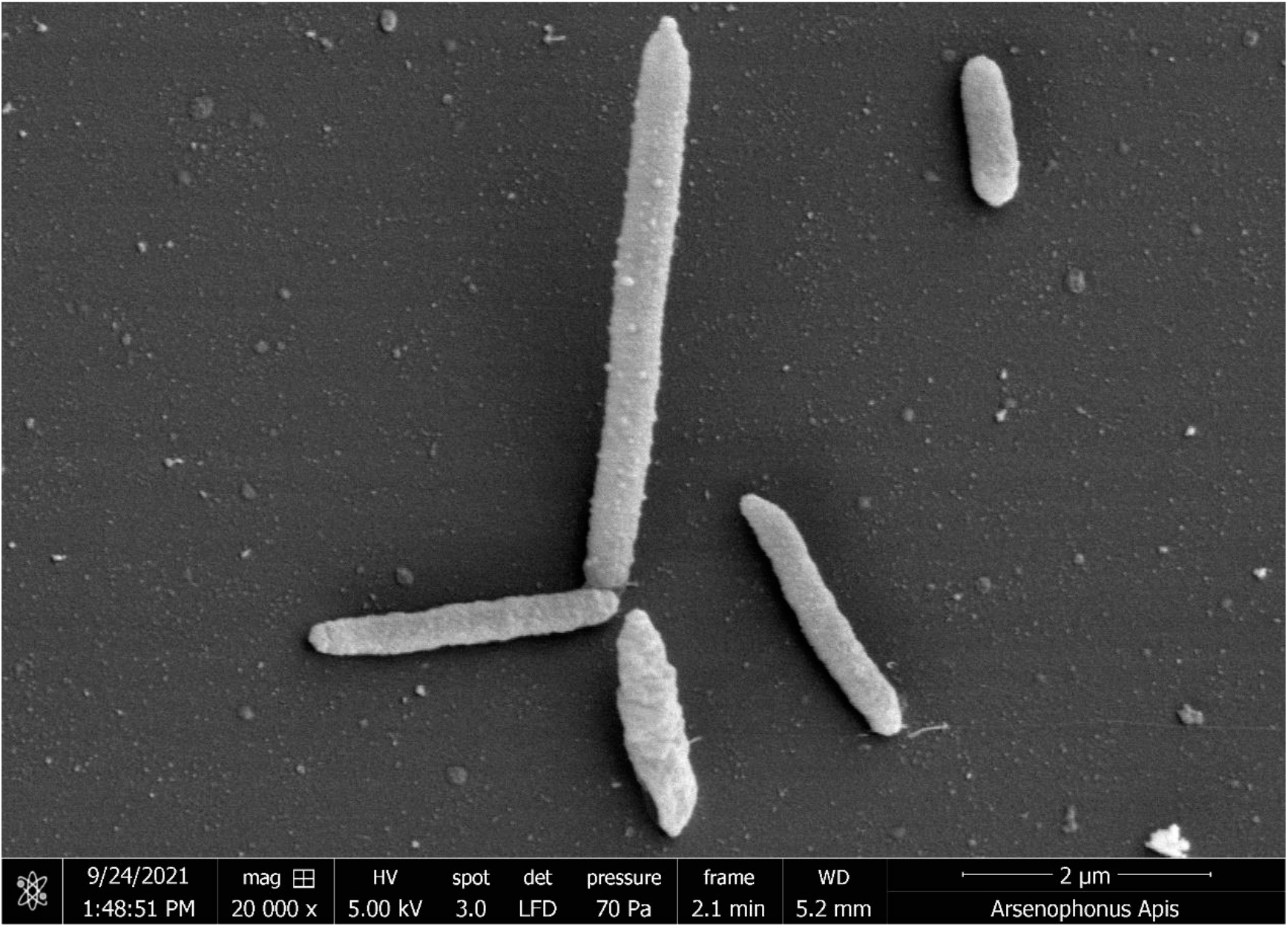
Scanning electron micrograph of *Arsenophonus apicola*. Scale bar = 2 μm

Phylogenetic analysis of the sequence of the 16S rRNA gene revealed the isolated microbe to lie within the genus *Arsenophonus*, most closely related to *A. nasoniae* (Figure 2).

**Figure 2.**
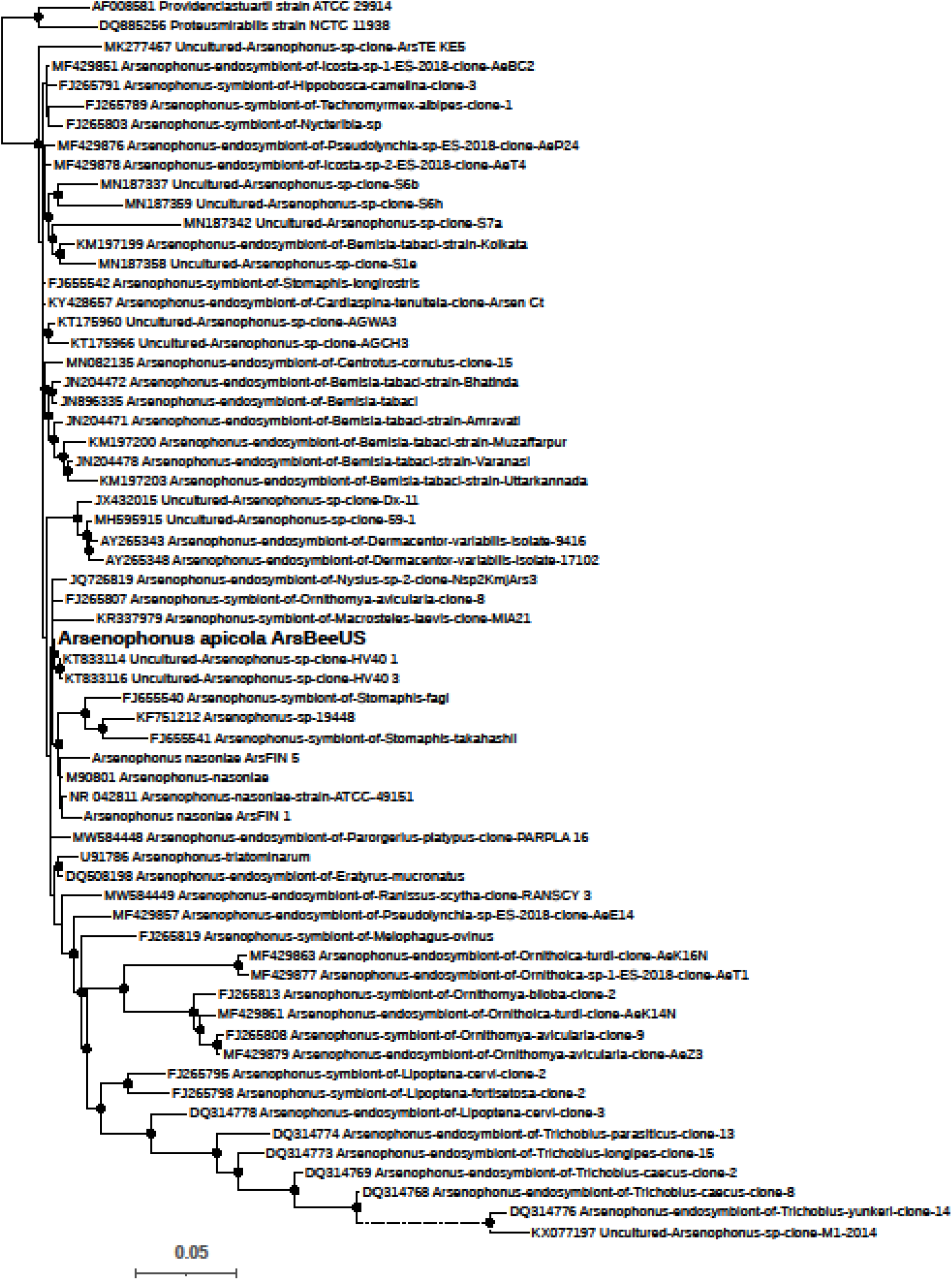
Phylogenetic position of *Arsenophonus apicola* in relation to other members of the genus Arsenophonus based on sequence of the 16S rRNA gene. Tree topology was inferred using maximum likelihood in IQ-TREE 2.1.4. Support values are based on 1000 ultrafast bootstrap replicates. The black circles on the nodes indicate support values greater than 80%. Dotted line indicates that the branch has been reduced by 50% for visualisation purposes.

### *In vitro* Growth requirements and *in vivo* growth

Biolog analysis indicated a restricted range of carbon utilization sources, although the range of carbon utilization sources extended beyond that of *A. nasoniae*, the type species (Table 1). The strain showed widespread resistance to external stressors and xenobiotics (Table 2).

**Table 1:** Utilization of carbon sources for growth by *A. apicola* with comparison to *A. nasoniae*. +++ strong growth, - growth absent.

|  | *A. apicola* | *A. nasoniae* |
| --- | --- | --- |
| *Dextrin* | - | - |
| *D-maltose* | - | - |
| *D-Trehalose* | - | - |
| *D-Cellulobiose* | - | - |
| *Gentiobiose* | - | - |
| *Sucrose* | - | - |
| *D-Turanose* | - | - |
| *Stachyose* | - | - |
| *D-raffinose* | - | - |
| *α-D-lactose* | - | - |
| *D-melibiose* | - | - |
| *B-Methyl-D-Glucoside* | - | - |
| *D-Salicin* | - | - |
| *N-Acetyl-D-Glucosamine* | +++ | +++ |
| *N-Acetyl-B-D-Mannosamine* | - | - |
| *N-Acetyl-D-Galactosamine* | - | - |
| *N-Acetyl Neuraminic Acid* | - | - |
| *α-D-Glucose* | +++ | +++ |
| *D-mannose* | +++ | +++ |
| *D-Fructose* | +++ | +++ |
| *D-Galactose* | - | - |
| *3-Methyl Glucose* | - | - |
| *D-Fucose* | - | - |
| *L-Fucose* | - | - |
| *L-Rhamnose* | - | - |
| *Inosine* | - | - |
| *D-sorbitol* | - | - |
| *D-mannitol* | - | - |
| *D-Arabitol* | - | - |
| *myo-inositol* | - | - |
| *Glycerol* | - | - |
| *D-Glucose-6-PO4* | +++ | +++ |
| *D-Fructose-6-PO4* | +++ | +++ |
| *D-Aspartic Acid* | - | - |
| *D-Serine* | - | - |
| *Gelatin* | - | - |
| *Glycyl-L-Proline* | - | - |
| *L-Alanine* | - | - |
| *L-Arginine* | - | - |
| *L-Aspartic Acid* | +++ | - |
| *L-Glutamic Acid* | ++ | ++ |
| *L-Histidine* | - | - |
| *L-Pyroglutamic Acid* | - | - |
| *L-Serine* | - | - |
| *Pectin* | - | - |
| *D-Galacturonic acid* | - | - |
| *L-Galactonic Acid Lactone* | - | - |
| *D-Gluconic Acid* | +++ | - |
| *D-Glucuronic Acid* | - | - |
| *Glucuronamide* | ++ | - |
| *Mucic Acid* | - | - |
| *Quinic Acid* | - | - |
| *D-Saccharic Acid* | - | - |
| *p-hydroxy phenyl acetic acid* | - | - |
| *Methyl Pyruvate* | +++ | ++ |
| *D-Lactic Acid Methyl Ester* | - | - |
| *L-Lactic acid* | ++ | - |
| *Citric Acid* | - | - |
| *α-Keto-Glutaric-Acid* | +++ | ++ |
| *D-Malic Acid* | +++ | - |
| *L-Malic Acid* | +++ | +++ |
| *Bromo-Succinic Acid* | +++ | +++ |
| *Tween-40* | - | - |
| *Gamma-Amino-Butyric acid* | - | - |
| *α-Hydroxy-Butyric Acid* | ++ | - |
| *B-Hydroxy-D-L-Butyric Acid* | - | - |
| *α-Keto-Butiric-Acid* | ++ | ++ |
| *Acetoacetic Acid* | ++ | - |
| *Propionic Acid* | - | - |
| *Acetic Acid* | - | - |
| *Formic Acid* | - | - |

**Table 2:** Impact of environmental and xenobiotic stress conditions on *A. apicola* growth on Biolog III plates compared to *A. nasoniae*. +++ = maintains full growth under condition stated - = no growth under condition stated.

|  | *A. apicola* | *A. nasoniae* |
| --- | --- | --- |
| *pH 6* | +++ | +++ |
| *pH 5* | ++ | ++ |
| *1% NaCl* | +++ | +++ |
| *4% NaCl* | +++ | +++ |
| *8% NaCl* | +++ | +++ |
| *1% Sodium lactate* | +++ | +++ |
| *Fusidic acid* | +++ | +++ |
| *D-Serine* | +++ | +++ |
| *Troleandomycin* | +++ | +++ |
| *Rifamycin SV* | +++ | +++ |
| *Minocycline* | +++ | +++ |
| *Lincomycin* | +++ | +++ |
| *Guanidine HCl* | +++ | +++ |
| *Niaproof 4* | - | ++ |
| *Vancomycin* | +++ | +++ |
| *Tetrazolium Violet* | - | ++ |
| *Tetrazolium Blue* | +++ | +++ |
| *Nalidixic Acid* | +++ | +++ |
| *Lithium Chloride* | +++ | +++ |
| *Potassium Tellurite* | +++ | +++ |
| *Aztreonam* | +++ | +++ |
| *Sodium Butyrate* | +++ | +++ |
| *Sodium Bromate* | - | - |

Substantial microbe growth within *A. mellifera* was observed in a three day period. At the whole bee level, a weak infection was observed one day post inoculation as evidenced from GFP fluorescence, but strong growth was evident after three days. Sham injected controls showed no GFP fluorescence (Figure 3; all bees imaged available at: https://doi.org/10.6084/m9.figshare.17912168.v1). 58.3% of microbe inoculated bees died between day 1 and day 3 (n=24) compared to 30.8% sham-injected controls (n=13).

**Figure 3:**
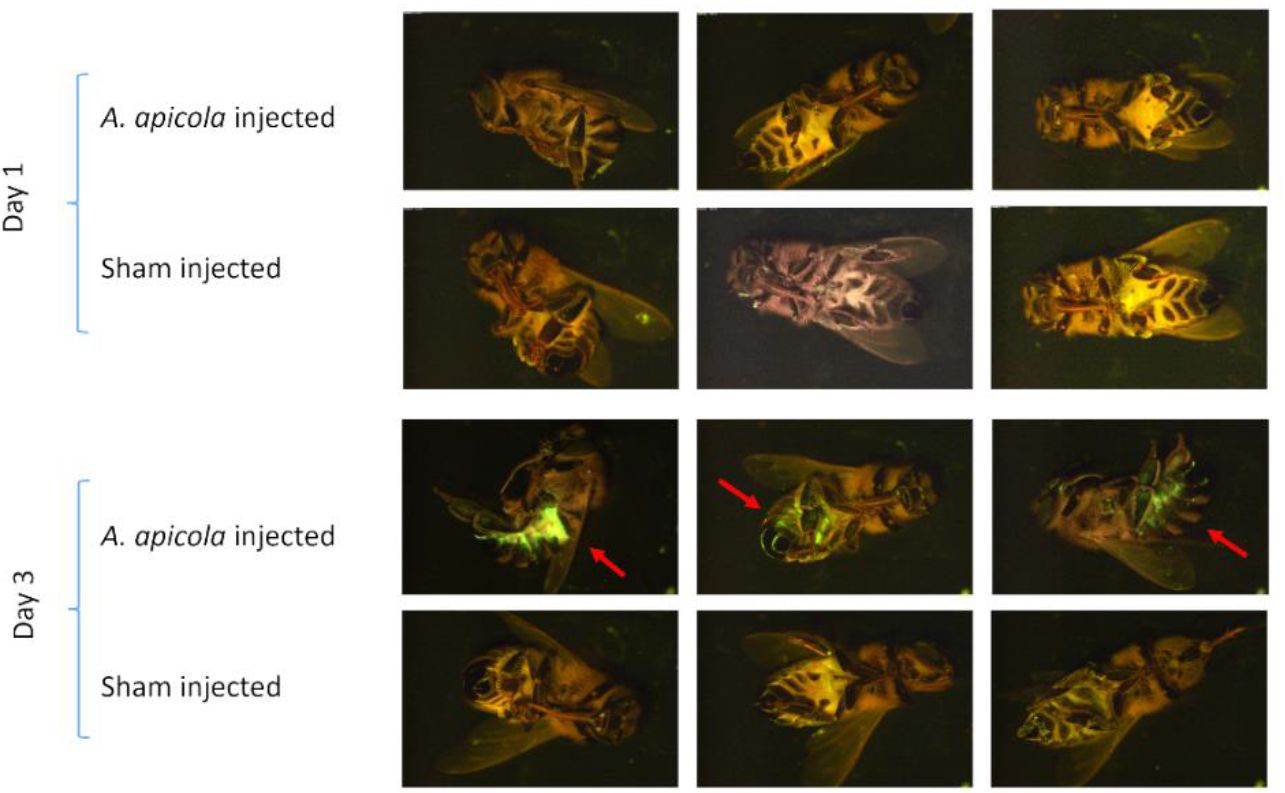
Epifluorescence images of *A. mellifera* worker bees one and three days after injection with *A. apicola* pOM1-*gfp* (top row) compared to sterile media-injected controls (bottom row).

### Genome sequence and assembly

The final assembly of *A. apicola* consists of a circular chromosome ~3.3 MB long with 6 circular plasmids with sizes range from ~90-30 KB (Table 3). The main chromosome has an average GC content of 37.6% and is predicted to contain 8 intact prophage regions and 1 incomplete prophage element. A further 3 intact prophage regions were identified within the plasmids. Genome annotation using the NCBI Prokaryotic Genome Annotation Pipeline identified a total of 3,190 protein coding sequences (CDSs) and an additional 242 pseudogenes accounting for a pseudogenization rate of circa 7.6%. Finally, 7 rRNA operons and 70 tRNA genes were identified in the main chromosome. The complete genome assembly has been submitted to the DDBJ/EMBL/GenBank database under the BioProject accession number PRJNA766690 (genome accession numbers CP084222 – CP084228).

**Table 3.**
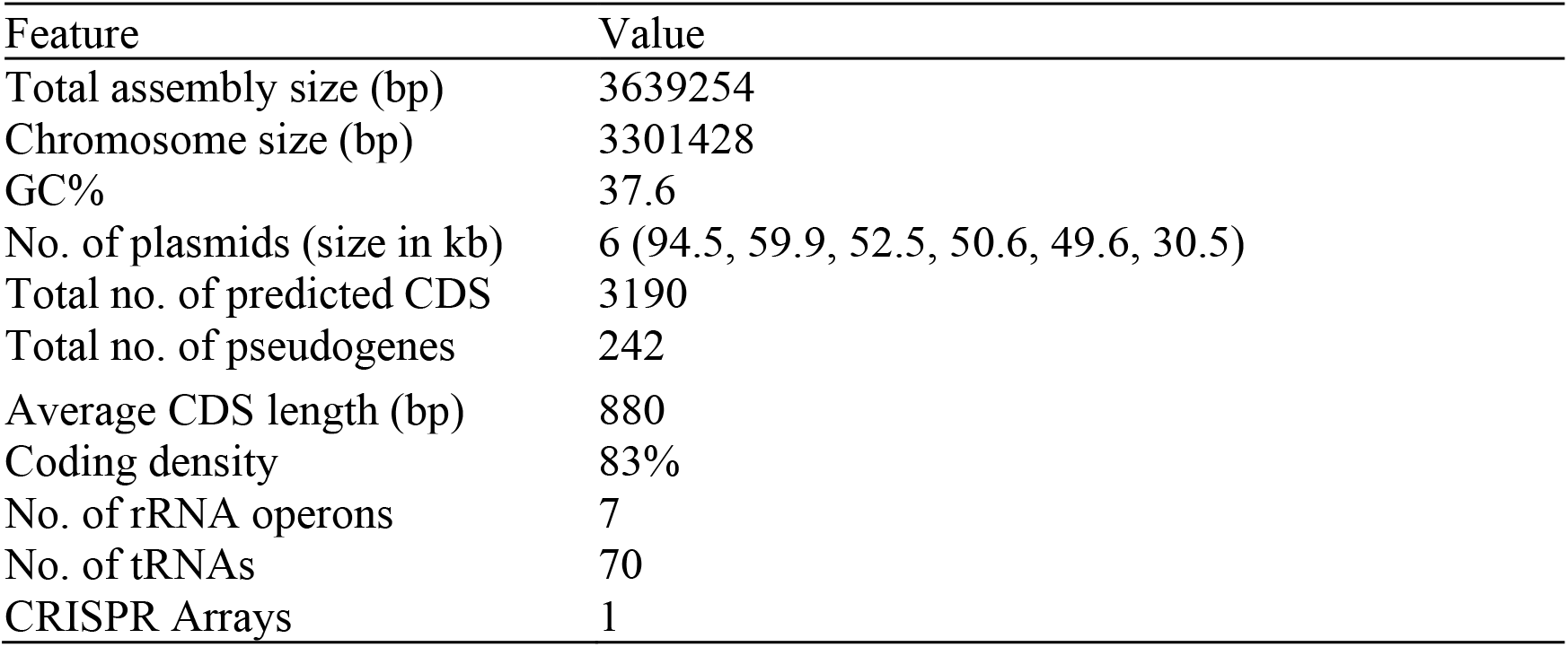
Genome features of *Arsenophonus apicola* strain ArsBeeUS.

| Feature | Value |
| --- | --- |
| Total assembly size (bp) | 3639254 |
| Chromosome size (bp) | 3301428 |
| GC% | 37.6 |
| No. of plasmids (size in kb) | 6 (94.5, 59.9, 52.5, 50.6, 49.6, 30.5) |
| Total no. of predicted CDS | 3190 |
| Total no. of pseudogenes | 242 |
| Average CDS length (bp) | 880 |
| Coding density | 83% |
| No. of rRNA operons | 7 |
| No. of tRNAs | 70 |
| CRISPR Arrays | 1 |

### Phylogenomic analysis, taxonomic classification and genomic analysis

Phylogenomic analysis using single-copy orthologues (Figure 4) placed *A. apicola* strain ArsBeeUS in the same clade with *Arsenophonus* previously reported from European honey bees (*Apis mellifera*) (20). The two strains share an ANI of >99% across the entire genome confirming they are the same species. The *Arsenophonus* from honey bees are closely related to *Arsenophonus nasoniae,* a male-killing reproductive parasite of *Nasonia vitripenis*. However, the ANI value between *A. nasoniae* and *A. apicola* is ~92% indicating they are separate species. The taxonomic novelty of the bee *Arsenophonus* clade was further confirmed using the GTDB taxonomic classifier.

**Figure 4.**
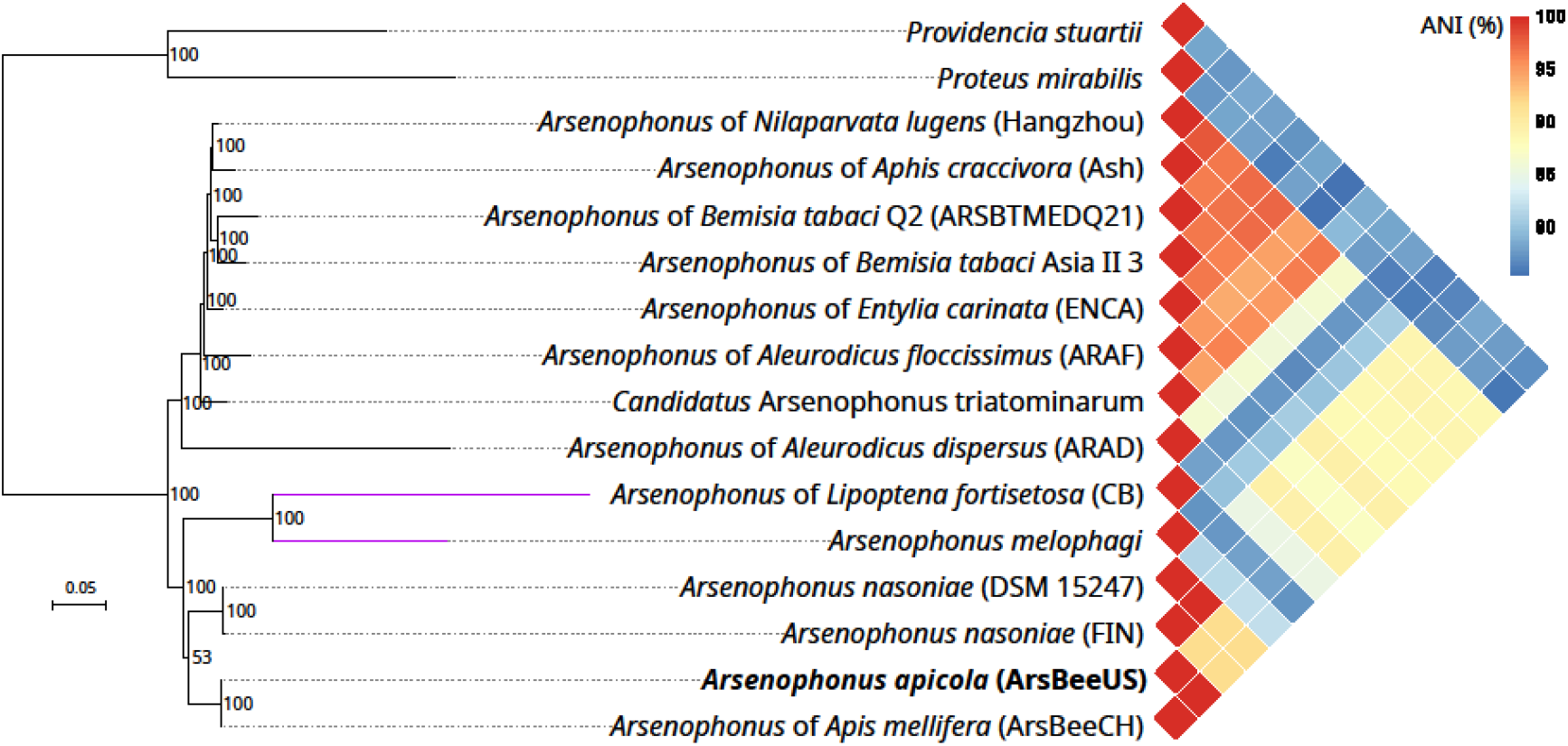
Whole genome phylogeny and average nucleotide identity (ANI). Phylogenetic relationships of *Arsenophonus* strains based on the concatenated analyses of 148 single copy core proteins. Species *Providencia stuartii* and *Proteus mirabilis* were used as outgroups. The relationships were inferred using IQ-TREE 2.1.4 and the maximum likelihood criterion. Support values are based on 1000 ultrafast bootstrap replicates. The purple branches have been reduced 50% for visualisation purposes. The triangular heatmap represents the pairwise ANI (%) values between the *Arsenophonus* genomes. ANI values were computed using the ANI/AAI-Matrix calculator from the enveomics toolbox collection.

The genome is predicted to encode type III secretion systems alongside a wide array of predicted secreted toxins including putative insecticidal toxins (annotation table: https://doi.org/10.6084/m9.figshare.17912198.v1), compatible with its association with insect hosts. Anti-SMASH predicted the presence of six genomic regions associated with the biosynthesis of secondary metabolites. Among them were two putative non-ribosomal peptide synthase regions (NRPS) and a siderophore biosynthesis cluster with similarity to the putrebactin/avaroferin biosynthetic gene cluster from *Xenorhabdus budapestensis*. Similar siderophores were identified previously as virulence factors in entomopathogenic bacteria (42). As we previously showed, the metabolic potential of the *Arsenophonus* from honey bees resembles that of *Arsenophonus nasoniae* (20). The genome encodes complete biosynthetic pathways for several B vitamins including biotin, riboflavin, folate and pyridoxine.

### Description of *Arsenophonus apicola, sp. nov*

*Arsenophonus apicola* [a.pi’co.la. L. fem. n. apis bee; L. suff. -cola from L. n. incola inhabitant, dweller; N.L. n. (nominative in apposition) apicola bee-dweller]. *Arsenophonus apicola* is a gram-negative rod-shaped bacterium, which grows optimally at 30°C in brain heart infusion (BHI) medium (Oxoid, UK) forming colonies within 120 h. Using Biolog GENIII plates, the tested ArsBeeUS strain of *A. apicola* utilized the following carbon sources: D-glucose, D-mannose, D-fructose, methyl pyruvate, D-gluconic acid, L-aspartic acid, D-glucose-6-phosphate, D-fructose-6-phosphate, N-acetyl-D-glucosamine, α-keto glutaric acid, D-malic acid, L-malic acid, bromo succinic acid, L-glutamic acid, glucuronamide, L-lactic acid, α-hydroxy butyric acid, α-keto butyric acid, acetoacetic acid. Growth was sustained at pH5 and 8% NaCl, and growth was not inhibited by 1% sodium lactate, fusidic acid, D-serine, troleandomycin, rifamycin SV, minocycline, lincomycin, vancomycin, nalidixic acid or potassium tellurite.

Bacteria from this species have been recorded from *Apis mellifera* insects from the USA and Europe.

*Arsenophonus apicola* forms a cluster with a variety of uncultured symbionts of insects as well as the previously cultured type species *A. nasoniae* Gherna.

The type strain ArsBeeUS (LMG 32504, CECT 30499) was isolated from *Apis mellifera* Dunn County, Wisconsin, USA, 44°57′10.2”N 91°53′13.2”W. The genome consists of a circular chromosome of size 3,301,428 bp and DNA G+C content is 37.6%, with 6 plasmids. The complete genome sequence of the type strain is available at Genbank accessions CP084222 – CP084228 and 16S rRNA sequence at OL469802.

## Acknowledgements

We are thankful to Alison Beckett for SEM Electron microscopy imaging and Andrea Ku for access to *A. mellifera* for experiments and training RC in handling bees. This work was funded by a BBSRC grant to GH (grant BB/S017534/1), funding from the European Union’s Horizon 2020 research and innovation program under Marie Skłodowska-Curie grant agreements 704382 (to C.L.F.), and a Microbiology Society Harry Smith Summer studentship (to RC).

